# Blocky proline/glutamine patterns in the SFPQ intrinsically disordered region dictate paraspeckle formation as a distinct membraneless organelle

**DOI:** 10.1101/2025.02.21.639608

**Authors:** Hiro Takakuwa, Takao Yoda, Tomohiro Yamazaki, Tetsuro Hirose

## Abstract

Membraneless organelles (MLOs) formed through phase separation play crucial roles in various cellular processes. Many MLOs remain spatially compartmentalized, avoiding fusion or engulfment. MLOs are formed by dynamic multivalent interactions, often mediated by proteins with intrinsically disordered regions (IDRs). However, the molecular principles behind how IDRs maintain MLO independence remain poorly understood. Here, we investigated the proline/glutamine (P/Q)-rich IDR of SFPQ, a protein identified as a key factor in segregating paraspeckles from nuclear speckles. Paraspeckle segregation analyses, using SFPQ mutants tethered to NEAT1_2 long noncoding RNA, revealed that P/Q residues within the SFPQ IDR, conserved from humans to zebrafish, are crucial for its segregation activity. Beyond amino acid composition, the blocky patterns of P/Q residues are required for the segregation from nuclear speckles. Among human IDRs exhibiting PQ-block patterns, BRD4 IDR shows strong sequence similarity to the SFPQ IDR, and exhibits comparable segregation activity. Molecular dynamics simulation suggests that the PQ-blocky patterns required for the paraspeckle segregation do not correlate with the IDR characteristics necessary for self-assembly. Thus, these data suggest that the PQ-blocky patterns in IDRs represent a previously uncharacterized property that contributes to MLO independence, possibly through a mechanism distinct from the conventional phase separation-promoting function of IDRs.

## Introduction

Membraneless organelles (MLOs) are mesoscopic compartments formed through phase separation and play critical roles in cellular processes by concentrating specific biomolecules (1–5). Many MLOs are located in the highly organized nuclear interchromatin spaces. Nuclear MLOs include nucleoli, Cajal bodies, nuclear speckles (NSs), and paraspeckles (PSs), which are involved in gene expression and the assembly of gene expression machinery (1, 4, 5). Growing evidence suggests that these MLOs are formed by dynamic multivalent interactions between proteins and/or nucleic acids (1, 2, 5). In proteins, intrinsically disordered regions (IDRs) often play a central role in driving phase separation, alongside multivalent domain-ligand interactions. IDRs are characterized by amino acid sequences enriched in a limited subset of residues and lacking a stable three-dimensional structure. Various studies have demonstrated that multivalent interactions between specific IDRs play crucial roles in driving phase separation and shaping the biophysical properties of condensates, particularly in highly crowded molecular environments (6–9). In addition to IDRs, RNAs play a critical role in phase separation (10–13).

Long noncoding RNAs (lncRNAs) are fundamental regulators of cellular processes. A subset of lncRNAs termed architectural RNAs (arcRNAs) function as the scaffolds of MLOs (14–16). NEAT1_2 lncRNA serves as an essential architectural scaffold of PSs (17–20). PSs were initially defined as the nuclear domains found in close proximity to NSs and are enriched with the characteristic DBHS (drosophila behavior human splicing) family proteins (21, 22). PSs play key roles in various physiological and pathological conditions including cancer (23–29). NEAT1 lncRNA has two isoforms, NEAT1_1 (3.7 kb) and NEAT1_2 (22.7 kb) through alternative 3′ end processing (30, 31). NEAT1_2, but not NEAT1_1, is indispensable for PS formation (30–32). More than 60 PS proteins (PSPs) are enriched in PSs, and some of these PSPs, such as SFPQ, NONO and FUS, are essential for the processes of PS formation (22, 30, 33–37). NONO, FUS, RBM14, and the SWI/SNF complex are essential for PS assembly, and the IDRs of RBM14 and FUS play a crucial role in this process (30, 31, 33, 38, 39). PSs exhibit spherical or cylindrical shapes and possess a core–shell internal organization, with PSPs showing core, shell, or patchy localization patterns (39, 40). This characteristic core–shell organization is formed through micellization, a type of phase separation, distinct from liquid-liquid phase separation (15, 41, 42).

Through extensive CRISPR/Cas9 genome editing of the NEAT1 gene, we have revealed that NEAT1_2 lncRNA contains multiple functional RNA domains, including 5′ RNA stability domain (0–1 kb), PS assembly domain (8–16.6 kb), and 3′ triple helix structure (31, 41). Among hundreds of NEAT1 deletion mutant cell lines, we found that the Mini-PSs, mutant PSs constructed by Mini-NEAT1 containing the PS assembly domain, the RNA stability domain (0–1 kb), and the triple helix, were present within NSs (43). In other words, the deleted regions (1–8 kb and 16.6–22.6 kb) are essential for PS segregation from NSs. These segregation domains recruit several PSPs, such as SFPQ, BRG1, and HNRNPF, to the shell of PSs, and their localization to the PS shell is essential for PS segregation from NSs (43). Among these, we focused on SFPQ and found that its segregation activity requires both the oligomerization domain and the proline (P) and glutamine (Q) residues within the IDR (43). Interestingly, the IDR of SFPQ exhibits segregation activity from NSs, independent of its role in promoting PS assembly (43). However, the molecular principles of SFPQ IDR that govern the PS segregation process–including its amino acid composition and arrangement–remain to be fully understood. Here, we investigated the molecular principles of the SFPQ IDR using various mutants and found that the enrichment of proline and glutamine, which is evolutionarily conserved, is critical for its activity. Furthermore, both the amino acid composition and the blocky pattern of these residues plays a significant role in its function. Based on the molecular principles identified in this study, we searched for IDRs with segregation activity similar to the SFPQ IDR. Finally, BRD4 was identified as a candidate protein exhibiting segregation activity. Notably, its P/Q-rich domain displayed an activity level comparable to that of the SFPQ IDR, suggesting the potential generality of this principle.

## Results

### Subdomains of the SFPQ IDR required for PS segregation from NSs

In our previous study, we employed the MS2 tethering system to investigate which paraspeckle proteins can induce the segregation of Mini-PSs from NSs and established a Mini-NEAT1 cell line with a 6×MS2 binding site at its 5′ end (Mini-NEAT1/6×MS2@8.2 kb) (Figure 1A, upper) (43). In this cell line, Mini-PSs were incorporated into NSs in the absence of protein tethering (Figure 1A, bottom left). Meanwhile, tethering SFPQ WT to the shell of Mini-PSs induced their segregation from NSs (Figure 1A, bottom center). Here, we generated a variety of SFPQ IDR mutants and tethered them to Mini-PSs to explore the molecular rules that are critical for its segregation activity (Figure 1A, bottom right). To narrow down the regions of SFPQ IDR that are critical for PS segregation, we first generated three SFPQ IDR partial deletion mutants (Δ33–104aa, Δ105–188aa and Δ189–265aa) fused with MS2 coat protein (MCP). The 33–104aa region is enriched with both proline and glutamine residues, while the 105–188aa and the 189–265aa regions are enriched with only proline (Figure 1B, Figure 1—figure supplement 1A). As shown in our previous study, tethering of SFPQ WT induced PS segregation from NSs, whereas the ΔIDR mutant exhibited a moderate reduction in this activity, and the ΔRRM2/NOPS mutant showed a marked decrease (Figure 1C, D, Figure 1—figure supplement 1B). The ΔIDR mutant exhibited the moderate reduction, as it loses the segregation function mediated by the IDR but retains the ability to interact with endogenous SFPQ, NONO, and PSPC1. In contrast, the ΔRRM2/NOPS mutant, defective in these interactions, exhibited the marked decrease (43). The Δ33–104aa and Δ105–188aa mutants exhibited reduced activity comparable to ΔIDR mutant (Figure 1C, D, Figure 1—figure supplement 1B). Meanwhile, the Δ189–265aa mutant exhibited activity similar to SFPQ WT (Figure 1C, D, Figure 1—figure supplement 1B). As negative controls, overexpression of these MCP-SFPQ mutant proteins in parental Mini-NEAT1 cells did not induce PS segregation from NSs (Figure 1C, D, Figure 1—figure supplement 1B). These results suggest that the 33–104aa and 105–188aa regions play a crucial role in the functionality of SFPQ through its IDR. Taken together, these data also support the significance of proline and glutamine residues within SFPQ IDR in the PS segregation process from NSs.

**Figure 1.**
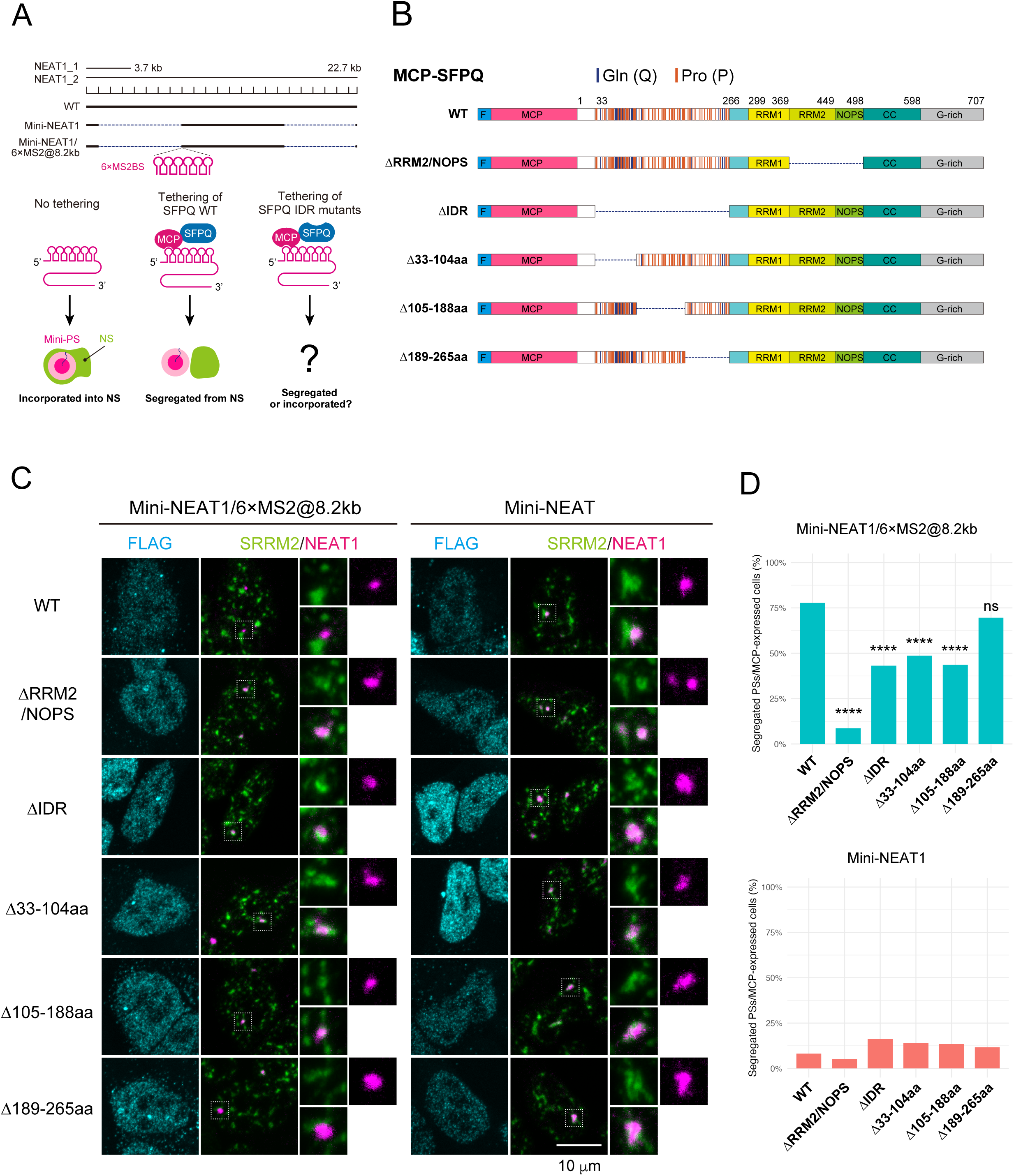
Subdomains of SFPQ IDR for PS Segregation from NSs. (A) Top: Schematic representation of NEAT1 wild-type (WT) and the NEAT1 mutants used in tethering experiments. The Mini-NEAT1 mutant features large deletions in the regions from 1–8 kb and 16.6–22.6 kb. The Mini-NEAT1/6×MS2@8.2kb mutant contains a 6×MS2 binding site inserted at the 8.2 kb position. Bottom: Schematic illustrating the MS2 tethering experiment. (B) Schematics of MCP-fused SFPQ WT and mutants used for tethering experiments. We referred to the region previously described as a PLD (prion-like domain), which was predicted by PLLAC (http://plaac.wi.mit.edu), as an IDR to maintain consistency in terminology. The positions of glutamine and glycine residues in the IDR of SFPQ are indicated with blue and orange lines, respectively. (C) Confocal observation of PSs (magenta) and NSs (green) with transfection of MCP-SFPQ WT and mutants into Mini-NEAT1/6×MS2@8.2Lkb (left) and Mini-NEAT1 (right) in the MG132 treatment conditions (5 μM for 6Lh). White boxes indicate the areas shown at a higher magnification. Scale bars, 10Lμm. (D) Quantification of PS segregation from NS. The data were collected from three independent experiments. p-values for comparisons with SFPQ WT were determined using two-tailed Fisher’s exact tests, with Holm correction for multiple comparisons. ****pL<L1.0 × 10^-4^. ns, not significant. Exact p-values are listed in the ‘Statistics and reproducibility’ section of the Methods. In Mini-NEAT1/6×MS2 cells expressing MCP-fused SFPQ variants: WT, n = 135; ΔRRM2/NOPS, n = 81; ΔIDR, n = 81; Δ33–104aa, n = 119; Δ105–188aa, n = 94; Δ189–265aa, n = 125. In Mini-NEAT1 cells expressing MCP-fused SFPQ variants: WT, n = 122; ΔRRM2/NOPS, n = 77; ΔIDR, n = 135; Δ33–104aa, n = 143; Δ105–188aa, n = 134; Δ189–265aa, n = 120.

### Amino acid composition of the SFPQ IDR is conserved in humans and zebrafish

IDRs often show poor sequence conservation, but the amino acid composition and specific physicochemical properties are conserved between species (9, 44, 45). Therefore, by examining the conservation of amino acid sequences across species, the amino acid composition that plays a critical role in IDR function can be identified (9, 45). We performed a multiple sequence alignment of SFPQ across nine vertebrate species, from humans to zebrafish. Consequently, it was revealed that the IDR of SFPQ is highly conserved from human to mouse (> 80%), while the sequence appears to be more variable from chicken to zebrafish (< 50%) (Figure 2A, Figure 2—figure supplement 1A, B). The IDR of zebrafish showed the greatest divergence compared to humans (Figure 2—figure supplement 1B). However, it was found that, similar to the human IDR, the zebrafish IDR is also predominantly enriched in proline and glutamine (Figure 2A, Figure 2—figure supplement 1C). To examine whether the IDR of zebrafish is functionally conserved, we generated a construct (MCP-hSFPQ-zIDR) in which the IDR of human SFPQ was swapped with that of zebrafish and performed the tethering assay (Figure 2B). The tethering of MCP-hSFPQ-zIDR rescued segregation defects fully to a level comparable to that of MCP-hSFPQ-WT, whereas the ΔIDR mutant did not rescue well (Figure 2C, D, Figure 2—figure supplement 1D). Based on these results, it is suggested that while the primary sequence conservation of the IDR of human and zebrafish SFPQ is low, the composition of amino acids necessary for its function is conserved.

**Figure 2.**
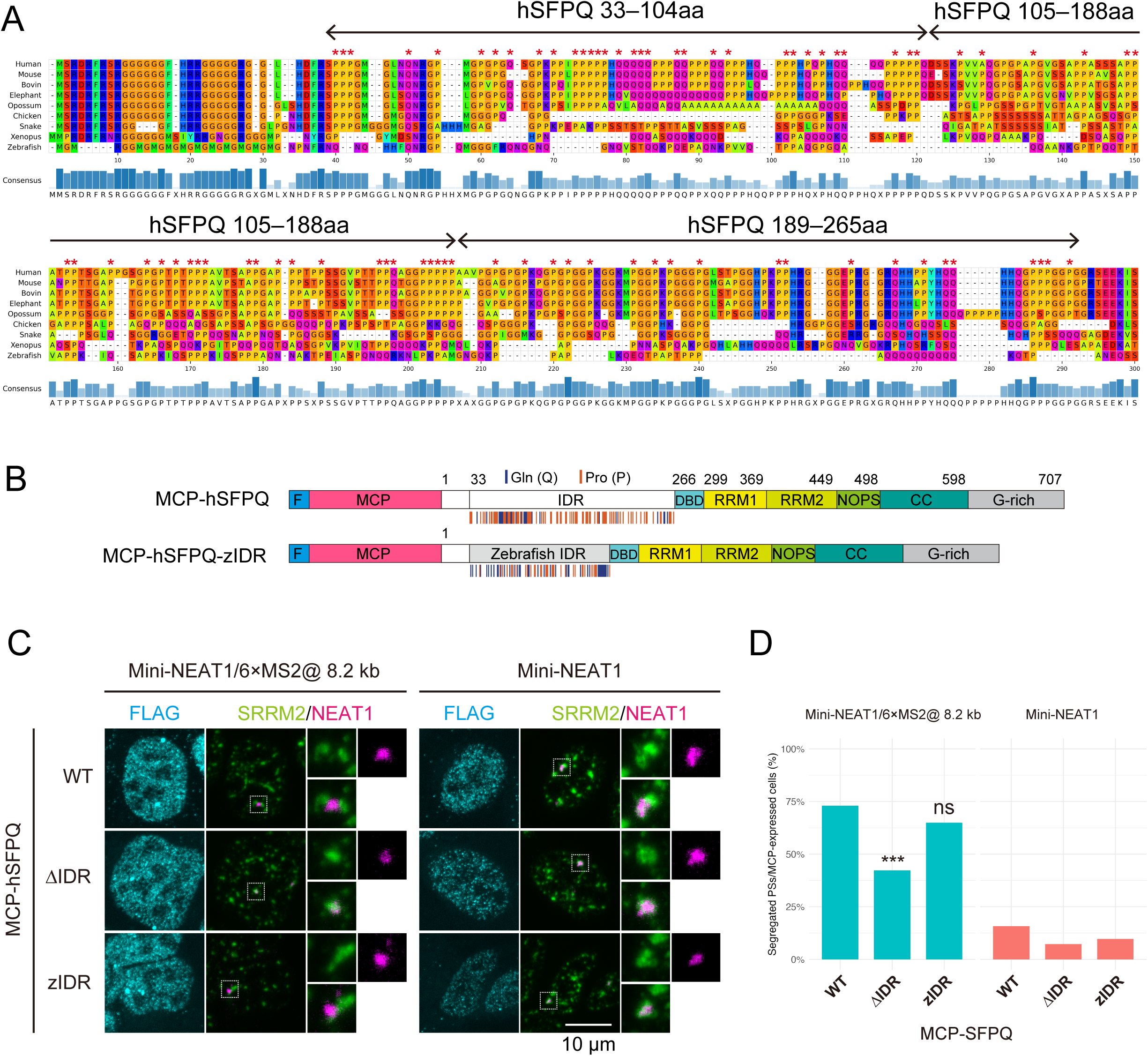
Function of SFPQ IDR in humans is conserved in zebrafish. (A) Multiple sequence alignment of SFPQ IDR from *Homo sapiens* (human), *Mus musculus* (mouse), *Bos taurus* (bovine), *Loxodonta africana* (elephant), *Monodelphis domestica* (opossum), *Gallus gallus* (chicken), *Pseudonaja textilis* (snake), *Xenopus laevis* (frog), and *Danio rerio* (zebrafish). The conserved proline and glutamine residues in 5 or more out of 9 species are highlighted by red asterisks on the top of the alignment. (B) Schematics of MCP-fused human SFPQ (MCP-hSFPQ) and MCP-fused human SFPQ with zebrafish IDR (MCP-hSFPQ-zIDR). (C) Confocal observation of PSs and NSs with transfection of MCP-hSFPQ WT and mutants into Mini-NEAT1/6×MS2@8.2kb (left) and Mini-NEAT1 (right) in the MG132 treatment conditions (5 μM for 6Lh). White boxes indicate the areas shown at a higher magnification. Scale bars, 10Lμm. (D) Quantification of PS segregation from NS. The data were collected from three independent experiments. p-values for comparisons with SFPQ WT were determined using two-tailed Fisher’s exact tests, with Holm correction for multiple comparisons. ***p < 1.0×10^-3^. ns, not significant. Exact p-values are listed in the ‘Statistics and reproducibility’ section of the Methods. In Mini-NEAT1/6×MS2 cells expressing MCP-fused SFPQ variants: WT, n = 100; ΔIDR, n = 78; zIDR, n = 77. In Mini-NEAT1 cells expressing MCP-fused SFPQ variants: WT, n = 120; ΔIDR, n = 96; zIDR, n = 103.

### Blocky proline and glutamine patterns in the IDR required for segregation activity

Previous studies have shown that in the IDRs of MED1 and Ki-67, the presence of patterned charge blocks plays a crucial role in phase separation (7, 46). These charge blocks have a stronger tendency to interact with each other compared to dispersed charge residues. To examine whether proline and glutamine are not only enriched but also exhibit sequence patterning features, we employed an unbiased computational analysis referred to as NARDINI+ (47, 48). Interestingly, within the 33–104aa region of human SFPQ, a blocky pattern of proline and glutamine–non-charged residues–is observed, as reflected by pro–pro, pol–pol, Q Patch, and P Patch metrics (Figure 3—figure supplement 1A). This prompted us to investigate whether the presence of patterned blocks of proline and glutamine residues in the IDR of SFPQ, along with their number, is critical for its function. To address this, we generated five mutants in which the 33–104aa region of the IDR was replaced with sequences composed exclusively of proline and glutamine, each with a different pattern (Figure 3A). In the tethering assay, the PPQQ (33–104aa) and P3Q3 (33–104aa) mutants exhibited segregation activity comparable to that of the WT, whereas the PQPQ (33–104aa), P12Q12 (33–104aa), and P36Q36 (33–104aa) mutants did not (Figure 3B, C, Figure 3—figure supplement 1B). These results suggest that a blocky pattern, characterized by alternating at least two proline and glutamine residues without excessively long continuous blocks, is crucial for the segregation activity of the SFPQ IDR. Next, we investigated whether the blocky pattern of proline and glutamine could compensate not only for the 33–104aa region but also for the 105–188aa region. We generated mutants in which the 33–188aa region was replaced with P3Q3 sequences of varying lengths and performed tethering assays (Figure 3A). Compared to the ΔIDR mutant, the P3Q3×3 (33–188aa) mutant showed little increase in the segregation activity, while the P3Q3×8 (33–188aa) and P3Q3×12 (33–188aa) mutants exhibited a moderate increase in this activity (Figure 3B, C, Figure 3—figure supplement 1B). These results suggest that the blocky pattern of proline and glutamine beyond a certain length is sufficient to compensate for the segregation function of the IDR in SFPQ.

**Figure 3.**
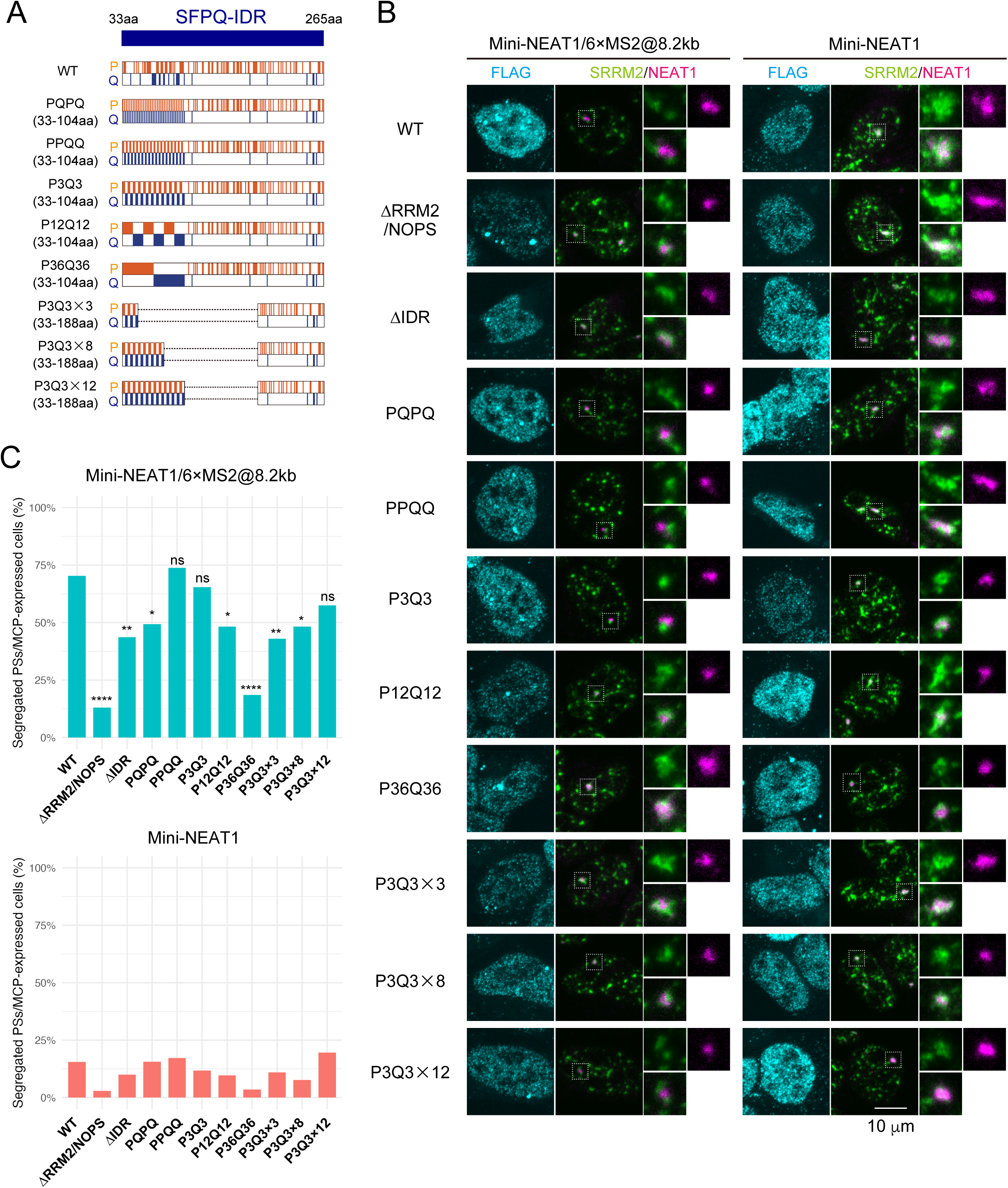
Blockiness of proline and glutamine in the SFPQ IDR. (A) Schematic representations illustrating the IDRs (33–265aa) for MCP-SFPQ WT and its mutants used in the tethering experiments. (B) Confocal observation of PSs and NSs with transfection of MCP-SFPQ WT and mutants into Mini-NEAT1/6×MS2@8.2kb and Mini-NEAT1 in the MG132 treatment conditions (5 μM for 6 h). White boxes indicate the areas shown at a higher magnification. Scale bars, 10Lμm. (C) Quantification of PS segregation from NS. The data were collected from three independent experiments. p-values for comparisons with SFPQ WT were determined using two-tailed Fisher’s exact tests, with Holm correction for multiple comparisons. ****p < 1.0×10^-4^, **p < 1.0×10^-2^, *p < 0.05. ns, not significant. Exact p-values are listed in the ‘Statistics and reproducibility’ section of the Methods. In Mini-NEAT1/6×MS2 cells expressing MCP-fused SFPQ variants: WT, n = 101; ΔRRM2/NOPS, n = 69; ΔPLD, n = 71; PQPQ, n = 79; PPQQ, n = 80; P3Q3, n = 84; P12Q12, n = 56; P36Q36, n = 92; P3Q3×3, n = 93; P3Q3×8, n = 85; P3Q3×12, n = 87. In Mini-NEAT1 cells expressing MCP-fused SFPQ variants: WT, n = 135; ΔRRM2/NOPS, n = 70; ΔPLD, n = 120; PQPQ, n = 115; PPQQ, n = 116; P3Q3, n = 94; P12Q12, n = 93; P36Q36, n = 116; P3Q3×3, n = 137; P3Q3×8, n = 104; P3Q3×12, n = 112.

### Coarse-grained MD simulations of P/Q-rich amino acid sequences

Our previous study demonstrated that the ability of SFPQ to rescue segregation defects is separable from its ability of paraspeckle assembly (43). However, the specific molecular actions of the SFPQ IDR in this process remain unclear. To investigate the role of the IDR in segregation activity, we conducted coarse-grained molecular dynamics (MD) simulations using a two-molecule system with WT (33–104aa), P3Q3 (33–104aa), P12Q12 (33–104aa), and P36Q36 (33–104aa) (Figure 4—figure supplement 1A). Upon measuring the temporal changes in the center-to-center distance between two molecules, the binding and dissociations of the two chains were observed for all variants (Figure 4A). Notably, it was also found that P12Q12 (33–104aa) and P36Q36 (33–104aa) exhibited more stable binding states compared to WT (33–104aa) and P3Q3 (33–104aa) (Figure 4A). The binding between these two molecules was primarily due to hydrophobic interactions between proline residues (Figure 4—figure supplement 1B). When calculating the spatial distribution of proline and glutamine from the center of geometry of the system, they turned out to be similarly distributed for both WT and P3Q3 (33–104aa) (Figure 4B). In contrast, P12Q12 (33–104aa) and P36Q36 (33–104aa) exhibited a configuration where proline residues were positioned more centrally, while glutamine residues were exposed to the solvent, as supported by their radial density profiles showing a central peak (∼0 Å) for prolines and a more peripheral distribution (∼10–16 Å) for glutamines (Figure 4B). These results suggest that P12Q12 (33–104aa) and P36Q36 (33–104aa) tend to aggregate more readily through proline compared to WT (33–104aa) and P3Q3 (33–104aa). It raises a possibility that the segregation activity of the SFPQ IDR may depend more on interactions with other molecules rather than relying on self-interactions to promote phase separation (see discussion).

**Figure 4.**
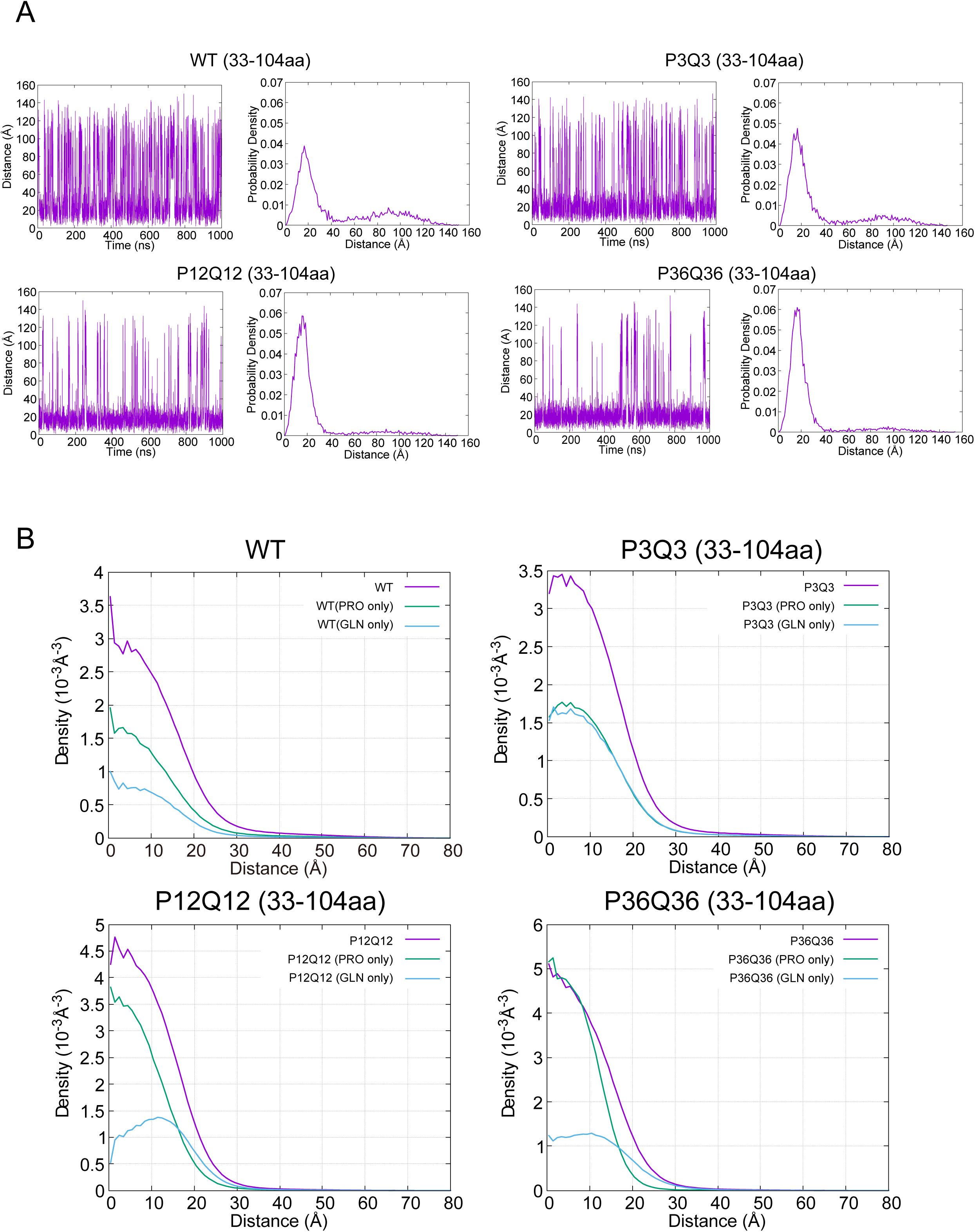
P3Q3 IDR is less likely to self-associate compared to P12Q12 and P36Q36 IDRs. (A) Time course of inter-center of geometry distance (left) and histogram of inter-center of geometry distance (right) between two molecules (B) The spatial distribution of amino acid residues in the two-molecule system. The distance from the center of geometry of the simulated system is plotted on the x-axis, with the volume density of proline and glutamine represented on the y-axis.

### Search for proteins with the segregation activity in IDR

Since the IDR with P/Q-rich blocky pattern exhibits segregation activity, we next explored whether proteins with similar IDR features have a similar function. To search for proteins that possess IDRs similar to that of SFPQ, we used information about IDR regions within the human proteome. We extracted data from MobiDB regarding IDR regions predicted to be disordered by AlphaFold2 (49). We then searched each IDR for 70 amino-acid sequences containing the highest total number of proline and glutamine residues, selecting this length because the SFPQ 33-104aa region spans 72 amino acids. We then calculated the P/Q-rich values (The number of P × the number of Q) to identify IDRs enriched in both proline and glutamine (Figure 5A, Figure 5—figure supplement 1A). Among the 13,971 IDRs, SFPQ exhibited the tenth-highest P/Q-rich value (Figure 5B, Figure 5—figure supplement 1A). To investigate the functional characteristics of the top 30 IDR proteins with the highest P/Q-rich values, we conducted a GO analysis (Figure 5C). In the biological process and molecular function categories, terms such as RNA metabolic process and transcription by RNA polymerase II were identified (Figure 5C). In the cellular component category, various GO terms were enriched, including chromatin, protein-DNA complexes, and notably, the SWI/SNF complex (Figure 5C). In our previous study, we identified BRG1/SMARCA4, a component of the SWI/SNF complex, as a protein with segregation activity, alongside SFPQ (43). In addition, ARID1A and SS18, other components of the SWI/SNF complex, were identified (Figure 5B). These results suggest that the IDRs of the SWI/SNF complex share common characteristics, which may contribute to their segregation activity. To further narrow down proteins with IDRs similar to that of SFPQ, we counted the occurrences of the blocky patterns (PPQQ or QQPP) and several other variations (Figure 5A, D, Figure 5—figure supplement 1B). Consequently, in addition to SFPQ, we found that the IDRs of KAT6A, FBXO11, SPRR5, and BRD4 were enriched in the blocky pattern by analyses using several block variations (Figure 5D, Figure 5—figure supplement 1B, C). Then, we selected the BRD4 IDR as a model and investigated whether its P/Q-rich IDR of BRD4 (from 742 to 1039 aa: Figure 5—figure supplement 1C) exhibits segregation activity from NSs, similar to that of SFPQ. We generated a chimera construct (MCP-SFPQ-BRD4 P/Q-rich), in which the IDR of human SFPQ was swapped with the P/Q-rich IDR of BRD4, and performed the tethering assay (Figure 5E). Tethering MCP-SFPQ-BRD4 P/Q-rich restored the reduced rescue activity observed in the ΔIDR mutant to a level comparable to that of MCP-hSFPQ-WT (Figure 5E, F, Figure 5—figure supplement 1D). These results suggest that the IDR of BRD4 exhibits activity comparable to that of the SFPQ IDR, indicating that P/Q-rich IDRs may have a functional role in biological systems.

**Figure 5.**
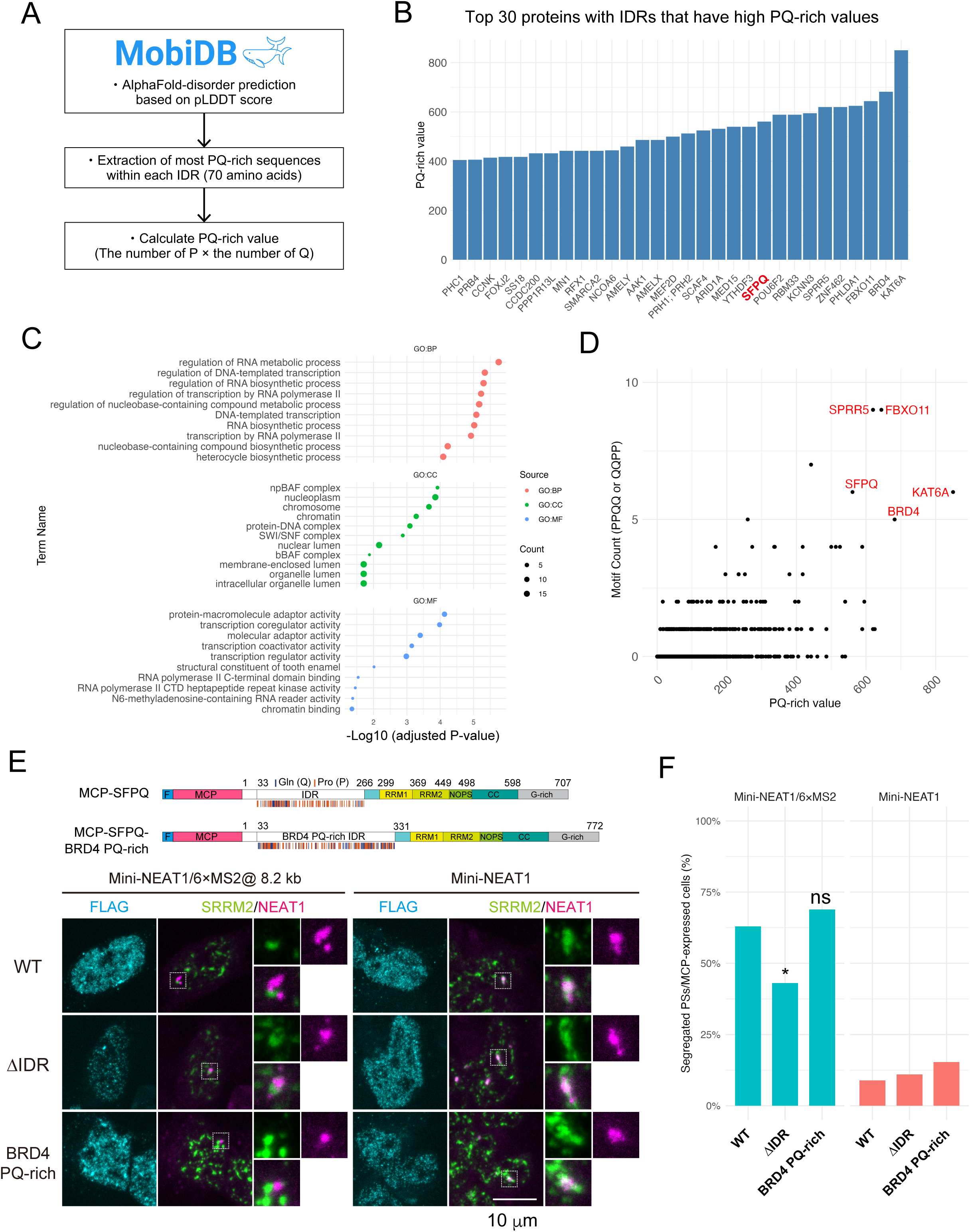
The IDR of BRD4 exhibits segregation activity. (A) Procedure for exploring IDRs with a blocky pattern of proline and glutamine. (B) Bar graph showing the top 30 proteins with IDRs that have high P/Q-rich values. (C) GO analysis of the top 30 proteins based on P/Q-rich values, showing the top 10 terms for GO Biological Process (GO:BP), Cellular Component (GO:CC), and Molecular Function (GO:MF), respectively. (D) Scatter plot with P/Q-rich values on the X-axis and the count of “PPQQ” or “QQPP” motifs on the Y-axis. The names of proteins with IDRs that have a P/Q-rich value of 500 or higher and a motif count of 5 or more are shown. (E) Top: Schematics of MCP-SFPQ WT and MCP-SFPQ with BRD4 P/Q-rich IDR (MCP-SFPQ-BRD4 P/Q-rich) used for tethering experiments. Bottom: Confocal observation of PSs and NSs with transfection of MCP-SFPQ WT and MCP-SFPQ-BRD4 P/Q-rich into Mini-NEAT1/6×MS2@8.2kb (left) and Mini-NEAT1 (right) in the MG132 treatment conditions (5 μM for 6 h). White boxes indicate the areas shown at a higher magnification. Scale bars, 10 μm. (F) Quantification of PS segregation from NS. The data were collected from three independent experiments. p-values for comparisons with SFPQ WT were determined using two-tailed Fisher’s exact tests, with Holm correction for multiple comparisons. *pL<L0.05. ns, not significant. Exact p-values are listed in the ‘Statistics and reproducibility’ section of the Methods. In Mini-NEAT1/6×MS2 cells expressing MCP-fused SFPQ variants: WT, n = 116; ΔIDR, n = 72; BRD4 PQ-rich (33–265aa), n = 90. In Mini-NEAT1 cells expressing MCP-fused SFPQ variants: WT, n = 101; ΔIDR, n = 100; BRD4 PQ-rich (33–265aa), n = 91.

## Discussion

We previously demonstrated that the IDR of SFPQ has two key functions: driving PS assembly and segregation of PSs from NSs. In this study, we uncovered a new molecular grammar within the SFPQ IDR–a P/Q-rich blocky pattern–which is critical for PS segregation. Additionally, by analyzing P/Q-rich proteins across the human proteome, we identified their association with gene expression pathways, offering valuable insights to advance future research on IDRs.

SFPQ is a multifunctional nuclear RNA-binding protein involved in various biological processes (37, 50–54). SFPQ possesses a modular domain structure, with many domains structurally and functionally characterized (55, 56); however, the role of its IDR remains largely unexplored in previous studies (57). Our study demonstrates that the 33–104aa and 105–188aa regions within the IDR play a crucial role in its segregation activity. While it remains unclear whether these regions have an equivalent function or play distinct roles, their unique amino acid composition offers intriguing insights. Notably, the 105–188aa region is slightly enriched with serine and threonine instead of glutamine (Figure 1—figure supplement 1A, Figure 3—figure supplement 1A). Considering serine, threonine, and glutamine are all non-charged and hydrophilic residues, the combination of hydrophobic proline with these uncharged, hydrophilic residues may be a crucial feature for segregation activity. Indeed, in chicken, snake, xenopus, and zebrafish, the region aligned with human 105–188aa contains more glutamine residues than in mammals (Figure 2A). In addition to the amino acid composition, we demonstrated that the blocky pattern of proline and glutamine is essential for the segregation activity of SFPQ (Figure 3). However, in zebrafish, the hydrophilic blocks consisting of glutamine in the IDR appear to be less conserved (Figure 2A, B). This might be explained by the presence of other uncharged, hydrophilic amino acids, such as threonine, serine, and asparagine, which are positioned near glutamine and could collectively form hydrophilic blocks (Figure 2A, B, Figure 2—figure supplement 1C). Additionally, focusing on the IDR in opossum, a part of the P/Q-rich region of human SFPQ IDR is substituted with a poly-alanine sequence (Figure 2A). Since paraspeckles are formed and segregated from nuclear speckles in opossum as in humans (58), the highly hydrophobic poly-alanine sequence may also possess a feature equivalent to that of the P/Q-rich sequence. The composition and arrangement of amino acids within IDRs vary across species, and by studying these variations, we may uncover previously unknown molecular grammars that govern phase behavior.

It is also important to note that there are technical limitations in the MD simulations. In this case, the simplification to a coarse-grained model resulted in structural details of the side chains not being fully captured. This is particularly relevant for proline, whose side chain forms a ring with the main chain, significantly limiting the possible conformational states it can adopt. Therefore, polyproline peptides adopt a secondary structure known as the polyproline-L (PPL) helix. The PPII helix is an extended (3.1 Å per residue compared to 1.5 Å in the α-helix) left-handed helix defined by the backbone torsion angle (φ,ψ) values of (−75°, 145°). Since PPL helices are shown to play important roles in protein-protein interactions and protein-nucleic acid interactions (59, 60), this structure may contribute to the segregation activity by the P/Q-rich IDR.

Further studies are required to fully elucidate the molecular mechanisms underlying the segregation of PSs from NSs through P/Q-rich IDRs. A recent study demonstrated that a blocky pattern of alanine, glycine, and glutamine residues in ARID1A/B, which are subunits of the SWI/SNF complex, is required for partner protein interactions rather than self-interactions (61). It has also been reported that ARID1B of the cBAF-type SWI/SNF complex interacts with NEAT1 lncRNA, and that the cBAF-type SWI/SNF complex includes BRG1/SMARCA4 and SS18, which are identified as proteins with IDRs showing high PQ-rich values (Figure 5B) (62). Since tethering the BRG1 induces the PS segregation from NSs independently of SFPQ, this function may operate as part of the cBAF-type SWI/SNF complex (43, 63). In addition, while the SPT6 WT IDR, which contains basic and acidic (charge) blocks, exhibited high affinity for MED1 IDR droplets in cells, the SPT6 IDR mutants with charge-to-P/Q-block substitutions showed high affinity for FUS IDR droplets but not for MED1 IDR droplets, suggesting that the presence of P/Q-blocks alters droplet affinity (8). The distinct functions of SFPQ IDR in PS assembly and PS segregation from NSs (43), along with the results of MD simulations, also support the possibility that blocky P/Q patterns function in the PS segregation by interacting with other partner molecules, such as proteins and nucleic acids. In line with this possibility, a recent in vitro phase separation study showed that the C-terminal IDR of SFPQ, rather than its N-terminal IDR, promotes homotypic interactions and facilitates droplet formation (57), suggesting that the N-terminal IDR induces PS segregation through a mechanism distinct from its LLPS capacity such as heteromeric interaction that may prevent NS interaction. However, this study does not address the identity of the heteromeric interaction partners. To elucidate this, it will be important to examine the potential existence of inhibitory factors that modulate NS interactions. Nevertheless, it remains challenging to determine whether the factors identified through interaction analyses are truly novel regulators that bind directly to IDRs and modulate their function. This represents a key question to be addressed in future studies.

Despite not being a major PS component, the BRD4 IDR exhibited PS-segregation activity comparable to that of the SFPQ IDR, raising the intriguing possibility that BRD4 may contribute to the independence of other MLOs, such as super-enhancer condensates where BRD4 resides (64). Thus, further investigation into the role of IDRs with P/Q-rich blocky patterns will provide new insights into the reciprocal interactions and phase immiscibility of multiple MLOs within cells.

## Materials and Methods

### Cell lines and cell culture

Human haploid cells (referred to as HAP1) were obtained from Horizon Discovery (Table S1). HAP1 cells were maintained in IMDM (Gibco) supplemented with 10% FBS (Sigma). The cells were grown at 37°C in a humidified incubator with 5% CO_2_. To induce NEAT1 expression, the cells were treated with 5 μM MG132 (Sigma-Aldrich) for 6 h.

### Plasmid construction

To construct plasmids expressing MCP-SFPQ IDR amino acid substitution mutant proteins, DNA fragments were introduced using the In-Fusion HD Cloning Kit (Takara Bio). To construct plasmids expressing MCP-SFPQ Δ33–104aa, Δ105–188aa, Δ189–265aa, the deletions were introduced by site-directed mutagenesis. Zebrafish IDR and BRD4 P/Q-rich region were amplified by PCR and replaced by In-Fusion reaction. SFPQ IDR mutants were synthesized by gene synthesis service (GenScript) (Table S1).

### Plasmid transfection

For MS2 tethering experiments, HAP1 cells were seeded in 12-well plates at a density of 2.0 × 10^5^ cells/well and cultured overnight. The cells were transfected with 1.0 μg of plasmid using 3 μl of TransIT-LT1 (Mirus), and the medium was changed 4 to 6 hours post-transfection. The cells were then cultured overnight. The cells were trypsinized and reseeded on coverslips (Matsunami, micro-cover glass, 18 mm round; thickness, 0.13–0.19 mm) at a density of 1.5 × 10^5^ cells/well. The cells were cultured for 24 hours, including the last 6 hours of MG132 treatment. Afterward, the cells were subjected to each experiment.

### RNA-FISH and immunofluorescence

RNA-FISH and immunofluorescence were performed as previously described (43). For MS2 tethering experiment, the cells were grown on coverslips and fixed with 4% paraformaldehyde/PBS at room temperature for 10 min. The cells were then washed with PBS, permeabilized with 0.5% Triton X-100/PBS for 5 min, and washed three times with PBS. Then, the coverslips were incubated with 1× blocking solution (Blocking Reagent [Roche] and PBST [PBS and 0.1% Tween 20]) at room temperature for 30 min. The coverslips were next incubated with primary antibodies in 1× blocking solution at 4L overnight, washed three times with PBST for 5 min, incubated with secondary antibodies in 1× blocking solution at room temperature for 2 h, and washed three times with PBST for 5 min. The primary antibodies used were anti-SRRM2 mouse monoclonal Ab (Sigma, S4045, 1:1000) and anti-DDDDK rabbit polyclonal Ab (PM020, MBL, 1:500). The secondary antibodies used were Goat anti-rabbit Alexa Fluor 405 (Invitrogen, A31556, 1:500) and Goat anti-mouse Alexa Fluor 488 (Invitrogen, A28175) (Table S1). After removing the PBST, RNA-FISH wash buffer (2× SSC and 10% formamide) was added and coverslips were incubated for 5 min. Then, 40 μl of hybridization solution (2× SSC, 100 mg/ml dextran sulfate, and 10% formamide) containing Quasar 570 Human NEAT1 custom probe sets (LGC Biosearch Technologies) at a final concentration of 125 nM were dropped onto the coverslips in a humidified chamber and incubated for 16 h in the dark. Next, the coverslips were washed with prewarmed RNA-FISH wash buffer at 37L for 30 min, and washed with 2× SSC at RT for 5 min. The coverslips were mounted with Prolong Diamond Antifade Mountant (Thermo Fisher Scientific). To visualize Mini-NEAT1 RNA, a mixture of the Quasar 570 Human NEAT1_5’ probe set and the Quasar 570 Human NEAT1_14.5–16.6 kb probe set (Table S2) was used. Images were acquired using a Nikon AX confocal microscope system with a Plan-Apochromat 60× oil immersion lens. Z-series images were captured at 0.5 μm intervals for 7 slices and processed using maximum intensity projection. Following a previous study, PSs were classified into “Segregated” and “Incorporated” types (43).

### Immunoblotting

Cells were lysed and denatured by adding SDS sample buffer and heating at 95°C for 5 min and then separated by SDS-PAGE. After electrophoresis, the proteins were transferred to a FluoroTrans W membrane (PALL) by electroblotting. The primary antibodies used were anti-FLAG mouse monoclonal Ab (Sigma, F3165, 1:5000) and anti-α-Tubulin mouse monoclonal Ab (Abcam, ab7291, 1:10000). The secondary antibody used was Mouse IgG HRP Linked Whole Ab (Cytiva, NA931, 1:5000) (Table S1).

### Multiple sequence alignment

Multiple sequence alignment was performed using Clustal Omega (https://www.ebi.ac.uk/jdispatcher/msa/clustalo). The resulting text files were visualized using pyMSAviz (https://github.com/moshi4/pyMSAviz).

### Analysis of IDRs using NARDINI+

The molecular grammar of IDRs was analyzed using the Grammars of IDRs using NARDINI+ (GIN) resource supported by Google Colab notebooks (https://colab.research.google.com/drive/1Lmb0pm5iFUOC4_ecBnFdmmcT0EOjLvfu#scrollTo=Sd7YgRpKAyY0) (47, 48). NARDINI+ algorithm yields a sequence specific z-score vector of 90 unique sequence features: 36 from NARDINI (65) and 54 from compositional analyses within the IDR sequence. To calculate patterning z-scores, the number of scrambles was set to 100,000. For patterning analysis, residues were grouped as follows: pol ≡ (S, T, N, Q, C, H), Aliphatic ≡ (I, L, M, V), pos ≡ (K, R), neg ≡ (E, D), Aromatic ≡ (F, W, Y), ala ≡ (A), pro ≡ (P), and gly ≡ (G). Z-scores less than zero imply the input sequence is more well-mixed than expected, whereas z-scores greater than zero imply the input sequence is blockier than expected.

For compositional features, human IDRome data analysed in the original study (48), were used to calculate compositional z-scores. Most compositional features were computed using the tool localCIDER (66) was utilized to calculate most compositional features. A “Patch” was defined as a region containing at least four occurrences of a given residue or two occurrences of RG with no more than two interruptions allowed.

### MD simulation

Coarse-grained molecular dynamics (CGMD) simulations of two-chain systems were carried out using version 1.7 of GENESIS atdyn after addition of codes needed for CGMD that are currently available in version 2.1 of GENESIS (67–69). The HPS CG model was employed for proteins (70). In the HPS model, one residue is represented by one bead each of which has amino-acid specific non-bonded interaction parameters to model hydrophobicity. The non-bonded interaction energy is defined by the following formula:

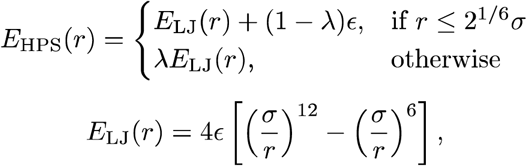

where *r* is the distance between two beads and ε = 0.2 kcal/mol. We used the original values of the amino-acid dependent hydrophobicity scale (λ) and the bead size (σ) (70). This non-bonded interaction was cutoff at 39 Å. The initial configuration of the simulated system contained two copies of a random-coil conformer that was generated by a 10-μs CG MD of the single-chain system of WT (33-104aa). Two chains were placed apart by 59 Å from each other in the cubic periodic-boundary box with 180 Å on each side. Net positive charge (+1) was assigned to Lys and Arg in WT (33-104aa), while the other beads were electrostatically neutral. Electrostatic interaction was modeled using a Coulombic term with Debye-Hückel electrostatic screening. Salt concentration and the cutoff distance for the electrostatic interaction were set to 0.15 M and 52 Å, respectively. Molecular dynamics simulations were performed for 2 μs at 300 K with NVT ensemble. The MD time step was 20 fs (velocity Verlet integrator). Temperature was maintained by Langevin thermostat with the friction parameter γ = 0.01 (ps^-1^). We used the latter half (1 μs) of the MD trajectories for analysis.

### Analysis of IDR database

Annotations on the IDRs from AlphaFold2 predicted structures were downloaded from MobiDB (version 5.0.0) (49). The unreviewed entries were excluded from the protein information obtained through UniProt ID mapping. IDRs longer than 70 amino acids were extracted, and within each IDR, the 70-amino-acid sequence with the highest total count of proline and glutamine residues was identified. To identify IDRs enriched in both proline and glutamine, the P/Q-rich value (the product of the number of proline and glutamine residues) was calculated. The number of ‘PPQQ’ and ‘QQPP’ motifs, as well as other similar patterns such as ‘PPPQQQ’ and ‘QQQPPP’ were counted to identify IDRs with a blocky pattern.

### Gene ontology analysis

Gene ontology analysis was conducted using g:Profiler (https://biit.cs.ut.ee/gprofiler/gost) with default settings. The top 10 terms for GO molecular function, GO cellular component, and GO biological process were included in Figure 5C.

### Statistics and reproducibility

All experiments were performed at least three times independently, with similar results obtained. No statistical method was used to predetermine the sample size. No data were excluded from the analyses. In some of the results from three independently performed experiments, the data from negative control and positive control are shared across experiments in Figure 1D, 2D, 3C, and 5F. A two-sample t-test with Bonferroni correction was employed in the analysis presented in Figure 5—figure supplement 1B and S2C. Fisher’s exact test and Holm correction were used for Figure 1D, 2D, 3C, and 5F. Exact p values were as follows: Figure 1D: p = 3.88×10^-24^ (ΔRRM2/NOPS), p = 1.21×10^-6^ (ΔIDR), p = 3.85×10^-6^ (Δ33-104aa), p = 6.81×10^-7^ (Δ105-188aa), p = 0.158 (Δ189-265aa); Figure 2D: p = 1.09×10^-4^ (ΔIDR), p = 0.255 (zIDR); Figure 3C: p = 4.14×10^-13^ (ΔRRM2NOPS), p = 3.81×10^-3^ (ΔIDR), p = 0.0275 (PQPQ), p = 1.00 (PPQQ), p = 1.00 (P3Q3), p = 0.0382 (P12Q12), p = 3.36×10^-12^ (P36Q36)), p = 0.0123 (P3Q3×3), p = 0.0159 (P3Q3×8), p = 0.278 (P3Q3×12); Figure 5F: p = 0.0204 (ΔIDR), p = 0.38 (BRD4 PQ-rich).

## Acknowledgments

The authors thank C. Fujikawa, and A. Kubota for technical support. A. Marshall, A.H. Fox, C.S. Bond and the members of the Hirose laboratory for valuable discussions. T. Yoda thanks Dr. Cheng Tan (RIKEN) and Dr. Yuji Sugita (RIKEN) for providing GENESIS atdyn source codes that enabled us CGMDs with the HPS model. This research was supported by supported by JST CREST grant no. JPMJCR20E6 (to T.H.), and JST FOREST program grant no JPMJFR2314 (to T. Yamazaki), AMED grant no. 21479280 (to T.H.), JSPS KAKENHI grant nos. 21H05282 (to T. Yoda), 22H02545 and 23H04258 (to T. Yamazaki), and 21H05276 and 24K21933 (to T.H.).

## Author contributions

H.T., T. Yamazaki, and T.H. conceived and designed this study. H.T. conducted the majority of the experiments. T. Yoda performed the MD simulations. H.T., T. Yamazaki, and T.H. wrote the manuscript.

## Declaration of interests

The authors declare no competing interests.

Figure 1—figure supplement 1. Supplemental data for Figure 1. (A) Enrichment or depletion of individual amino acids in 33–104aa (blue), 105–188aa (orange) and 189–265aa (green) of SFPQ IDR, as analyzed using Composition Profiler (http://www.cprofiler.org/). The SwissProt 51 dataset was used as the background. p values were determined by using a two-sample t-test, with Bonferroni correction for multiple comparisons. If the statistical test showed no significant difference (*p* > 0.05), it is not specified in the figure. (B) Immunoblotting of Flag-tagged MCP-SFPQ WT and mutants in the Mini-NEAT1 and Mini-NEAT1/6×MS2@8.2kb cells (Figure 1B–D). α-Tubulin is a loading control. Molecular weight markers (MW) are shown on the right.

Figure 2 —figure supplement 1. The central and C-terminal regions of SFPQ are relatively well conserved across species. (A) Multiple sequence alignment of full-length SFPQ from *Homo sapiens* (human), *Mus musculus* (mouse), *Bos taurus* (bovine), *Loxodonta africana* (elephant), *Monodelphis domestica* (opossum), *Gallus gallus* (chicken), *Pseudonaja textilis* (snake), *Xenopus laevis* (frog), and *Danio rerio* (zebrafish). (B) Guide tree of SFPQ IDRs from human to zebrafish. (C) Enrichment or depletion of individual amino acids in human SFPQ IDR (blue) and zebrafish SFPQ IDR (orange), as analyzed using Composition Profiler. The IDR region of zebrafish was defined as the corresponding region to the human SFPQ IDR in the alignment shown in Figure 2A. The SwissProt 51 dataset was used as the background. p values were determined by using a two-sample t-test, with Bonferroni correction for multiple comparisons. If the statistical test showed no significant difference (*p* > 0.05), it is not specified in the figure. (D) Immunoblotting of Flag-tagged MCP-SFPQ WT and mutants in the Mini-NEAT1 and Mini-NEAT1/6×MS2@8.2kb cells (Figure 2B–D). α-Tubulin is a loading control. Molecular weight markers (MW) are shown on the right.

Figure 3 —figure supplement 1. Supplemental data for Figure 3. (A) Compositional and patterning features of each subdomain within the SFPQ IDR, as analysed using the NARDINI+ algorithm. Only features with an absolute Z-score ≧ 1 in at least one subdomain are shown. (B) Immunoblotting of Flag-tagged MCP-SFPQ WT and mutants in the Mini-NEAT1 and Mini-NEAT1/6×MS2@8.2kb cells (Figure 3A–C). α-Tubulin is a loading control. Molecular weight markers (MW) are shown on the right.

Figure 4 —figure supplement 1. Supplemental data for Figure 4. (A) Final structure in the two-molecule system. (B) Intermolecular residue contacts in the two-molecule system. The quantity proportional to the probability of the residue distance being ≤ 7.0 Å is displayed on a color map.

Figure 5 —figure supplement 1. Supplemental data for Figure 5. BRD4 has IDR with a blocky pattern of proline and glutamine. (A) Table of the top 10 proteins based on P/Q-rich values. From left to right: Protein name, P/Q-rich sequence (70 amino acids), P/Q-rich values (The number of P × the number of Q), and Motif counts (“PPQQ” and “QQPP”). (B) Left: Scatter plot showing P/Q-rich values on the X-axis and the count of “PPXQQ” or “QQXPP” motifs on the Y-axis. Protein names are displayed for IDRs with a P/Q-rich value of 500 or higher and a motif count of at least 5. Right: Scatter plot with P/Q-rich values on the X-axis and the count of “PPPQQQ” or “QQQPPP” motifs on the Y-axis. Protein names are shown for IDRs with a P/Q-rich value of 500 or higher and a motif count of at least 2. (C) A diagram showing the positions of “PPQQ” and “QQPP” motifs in each protein. In Figure 5E and F, the region from 742 to 1039 aa of BRD4 is defined as the P/Q-rich IDR. (D) Immunoblotting of Flag-tagged MCP-SFPQ WT and mutants in the Mini-NEAT1 and Mini-NEAT1/6×MS2@8.2kb cells (Figure 5E, F). α-Tubulin is a loading control. Molecular weight markers (MW) are shown on the right.

