## Supplemental Figure S1, S2, S3, S4, S5, Supplemental Table S1, S2 for "Blocky proline/glutamine patterns in the SFPQ intrinsically disordered region dictate paraspeckle formation as a distinct membraneless organelle"

### Figure 1—figure supplement 1

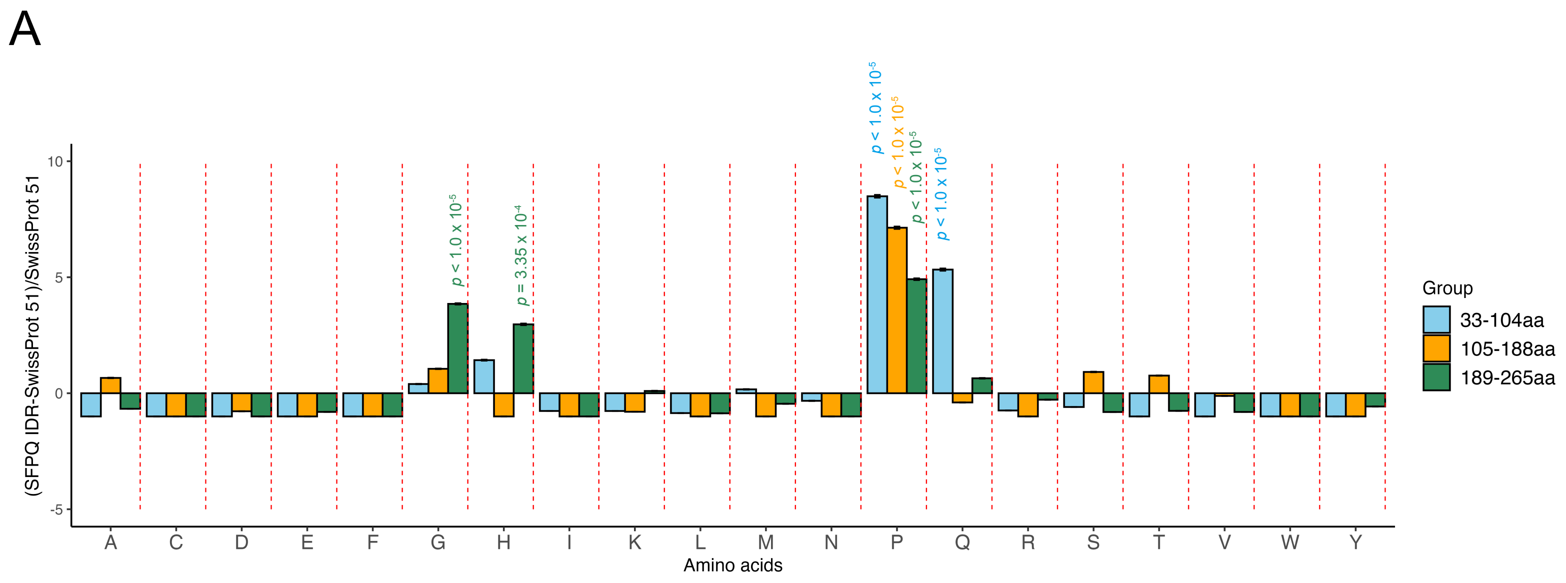


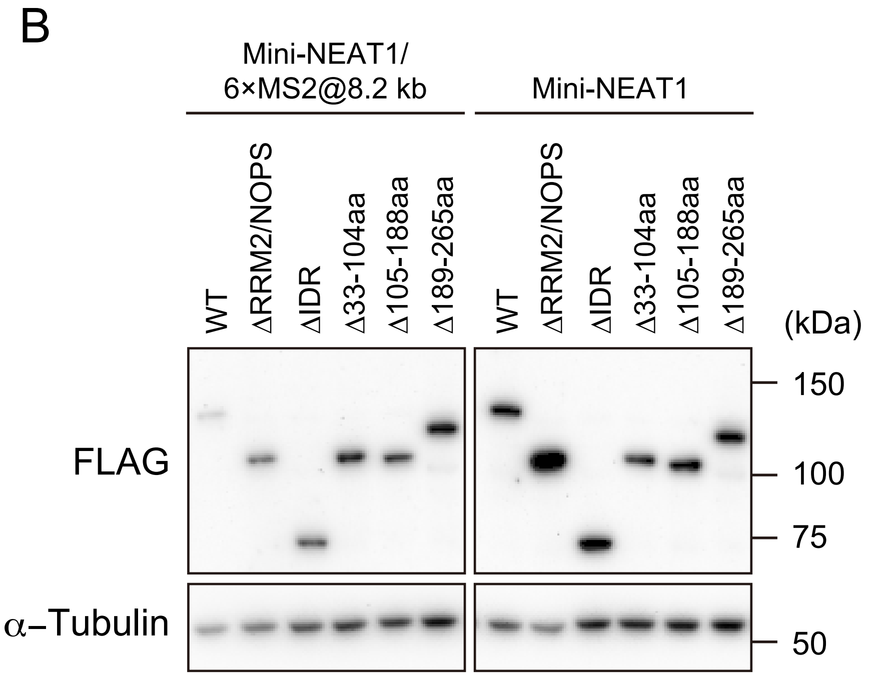

### Figure 2—figure supplement 1

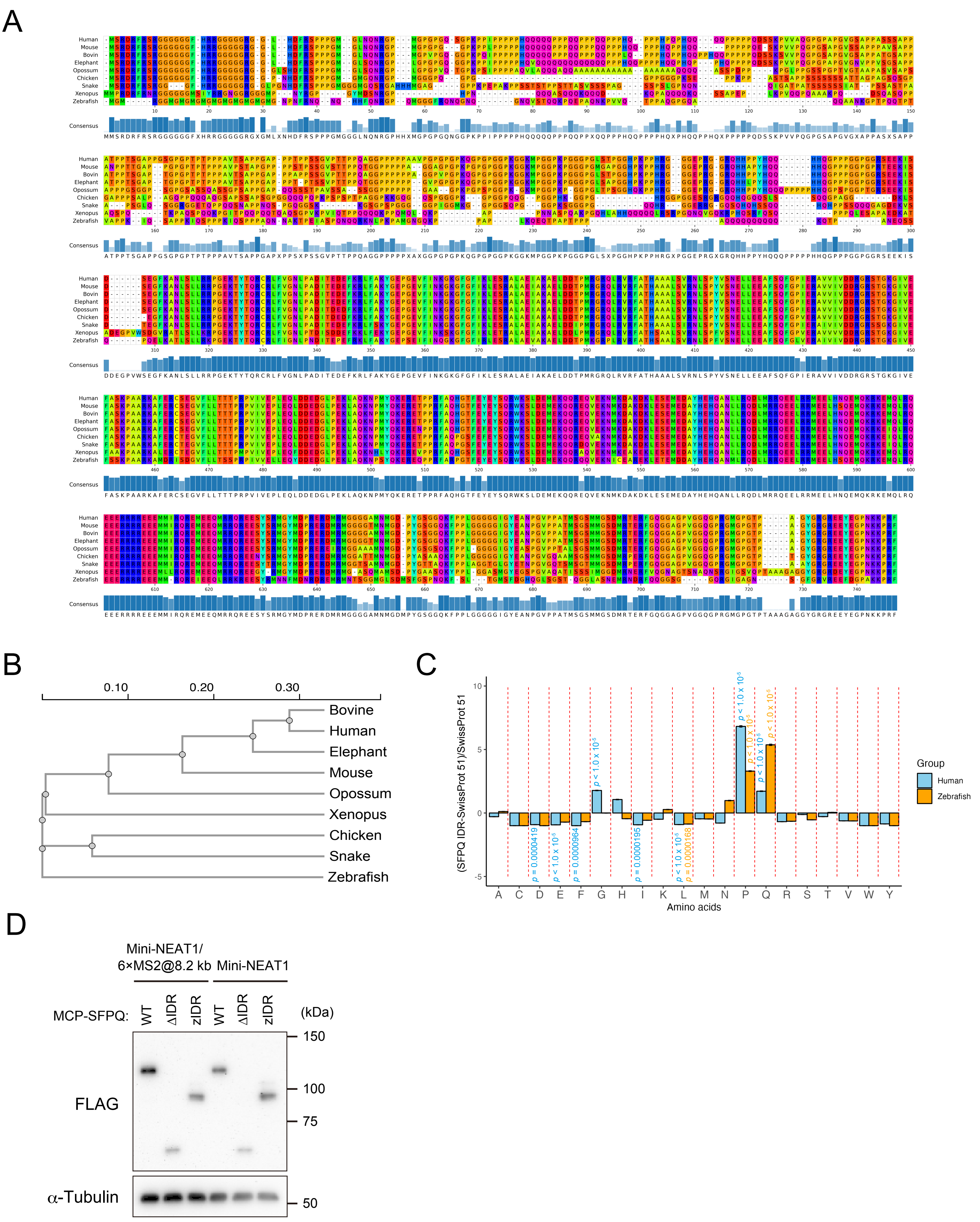

### Figure 3—figure supplement 1

A

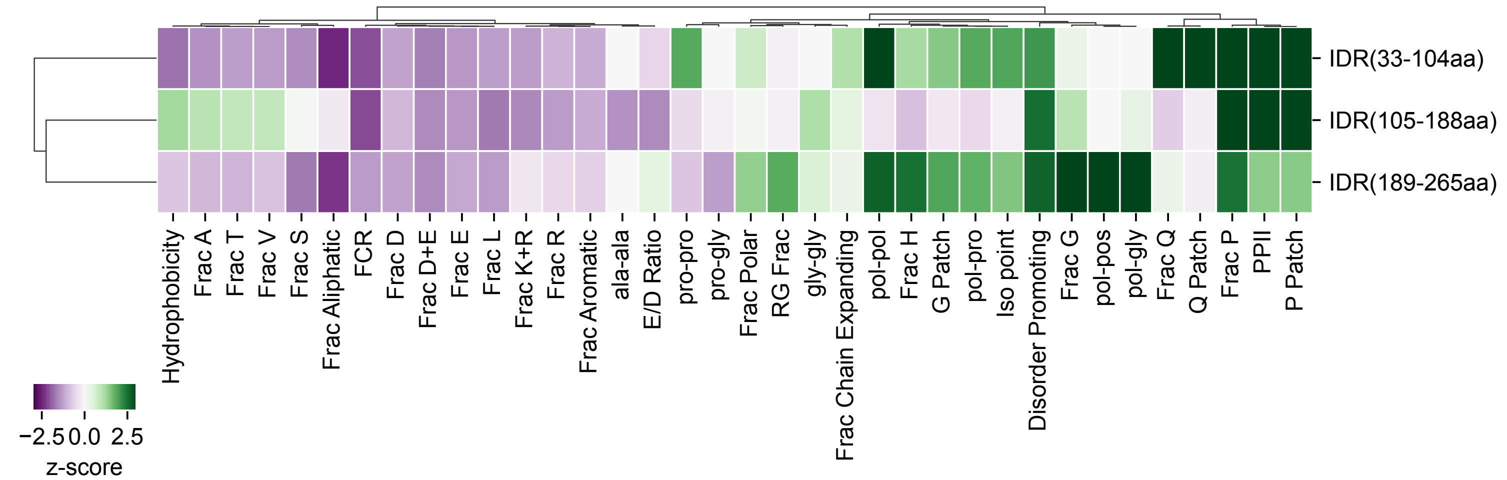


| | | | |
|---|---|---|---|
| pro: P | pol: S, T, N, Q, C, H | Aliphatic: A, I, L, M, V | PPII (polyproline-II propensity) |
| ala: A | pos: K, R | Aromatic: F, W, Y | Iso point (isoelectric point) |
| gly: G | | FCR: D, E, K, R | |

B

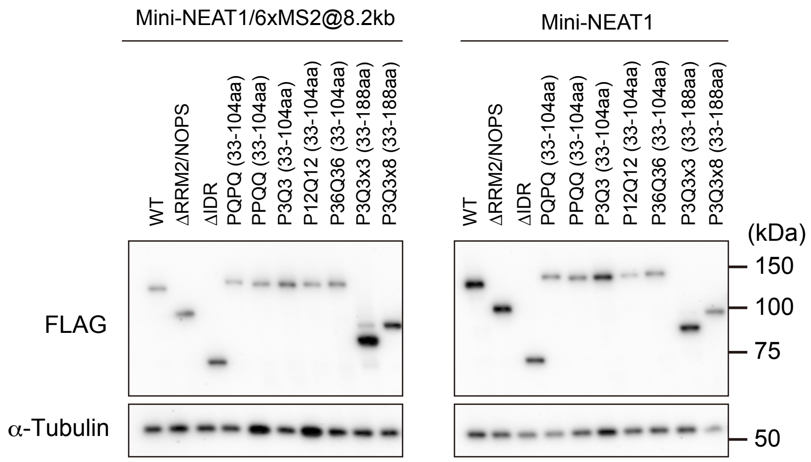

Figure 4—figure supplement 1

A

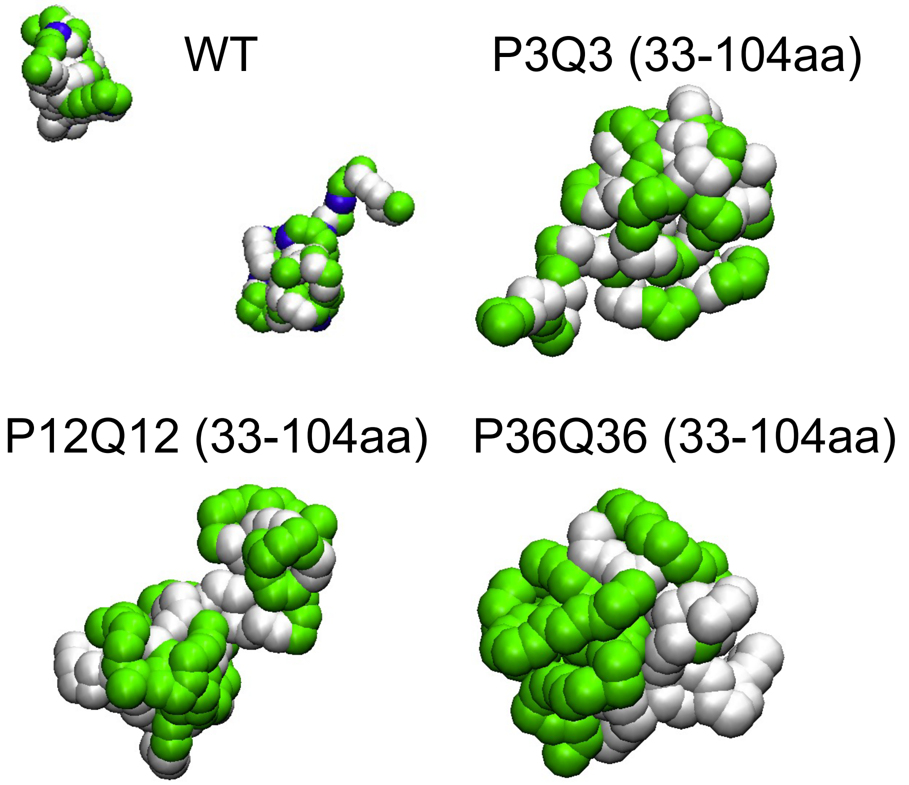


B

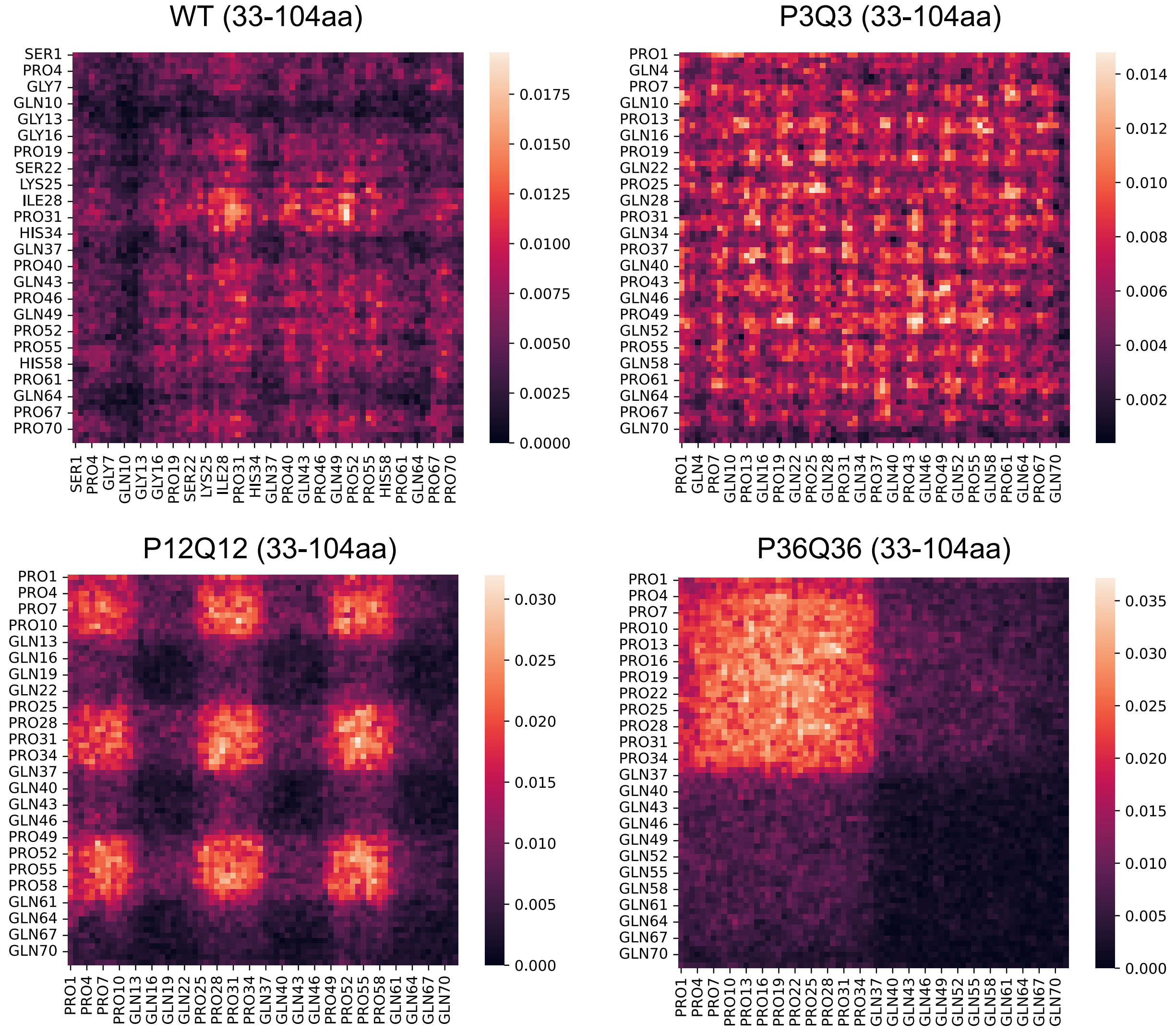

### Figure 5—figure supplement 1

A

| | Gene.Names..primary. | PQ_rich_sequence | PQ_rich_value | Motif_Count |
|---|---|---|---|---|
| 1 | KAT6A | PQSCVVERPPSNQQQQPPPPPPQQPQPPPPQPQPAPQPPPPQQQPQQQPQPQPQQPPPPPPPQQQPPLSQ | 850 | 6 |
| 2 | BRD4 | PSVQQQLQQQPPPPPPPQPQPPPQQQHQPPPRPVHLQPMQFSTHIQQPPPPQGQQPPHPPPGQQPPPPQP | 682 | 5 |
| 3 | FBXO11 | MNSVRAANRRPRRVSRPRPVQQQQQQPPQQPPPQPPQQQPPQQQPPPPPQQQQQQQPPPPPPPPPPLPQE | 644 | 9 |
| 4 | PHLDA1 | QKQQHLVQQQPPSQPQPQPQLQPQPQPQPQPQPQPQSQPQPQPQPKPQPQQLHPYPHPHPHPHSHPHSHP | 625 | 1 |
| 5 | SPRR5 | PPPQQRCPPPQQCCPPPQQCCPPPQQCCPPPQQCCPPPQQCCPPPQQCCPPPQQYCPPPQQTKQPCQPPP | 620 | 9 |
| 6 | ZNF462 | PLQQQQPPQPPPPPPPPPPPSQPQPLQQPQPPQLQPPHQVPPQPQTQPPPTQQPQPPTQAAPLHPYKCTMC | 620 | 1 |
| 7 | KCNN3 | EQQQQQQQQQQQQPPPPAPPAAPQQPLGPSLQPQPPQLQQQQQQQQQQQQQQPPHPLSQLAQLQSQPVHP | 595 | 2 |
| 8 | POU6F2 | QQLQLQLQQQQQQQQQQPPPSTNQHPQPAPQAPSQSQQQPLQPTPPQQPPPASQQPPAPTSQLQQAPQPQ | 589 | 4 |
| 9 | RBM33 | PGPGQPFLPTHTQPNLQGPLHPPLPPPHQPQPQQPQQQPPPQHQPPHQPPHQPPPQHQPPPQHPPQHPPQ | 589 | 1 |
| 10 | SFPQ | PPPGMGLNQNRGPMGPGPGQSGPKPPIPPPPPHQQQQPPPQQPPPQQPPPHQPPPHPQPHQQQQPPPPP | 561 | 6 |

B

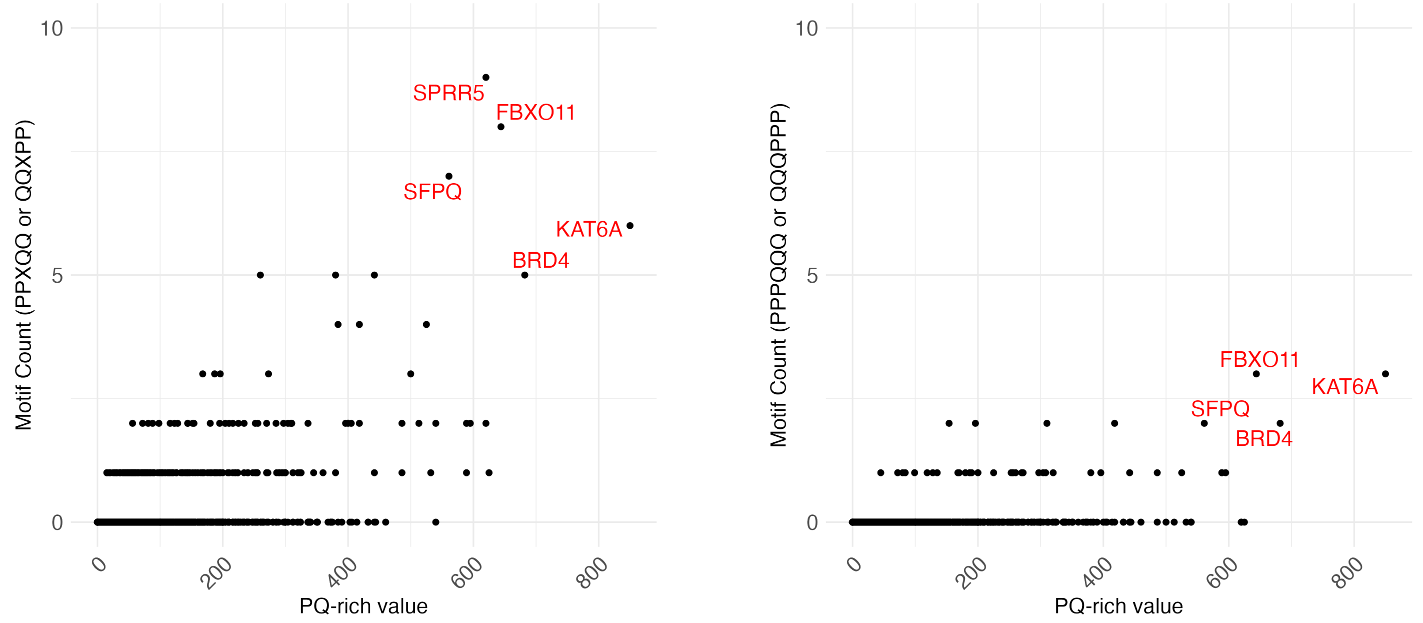


C

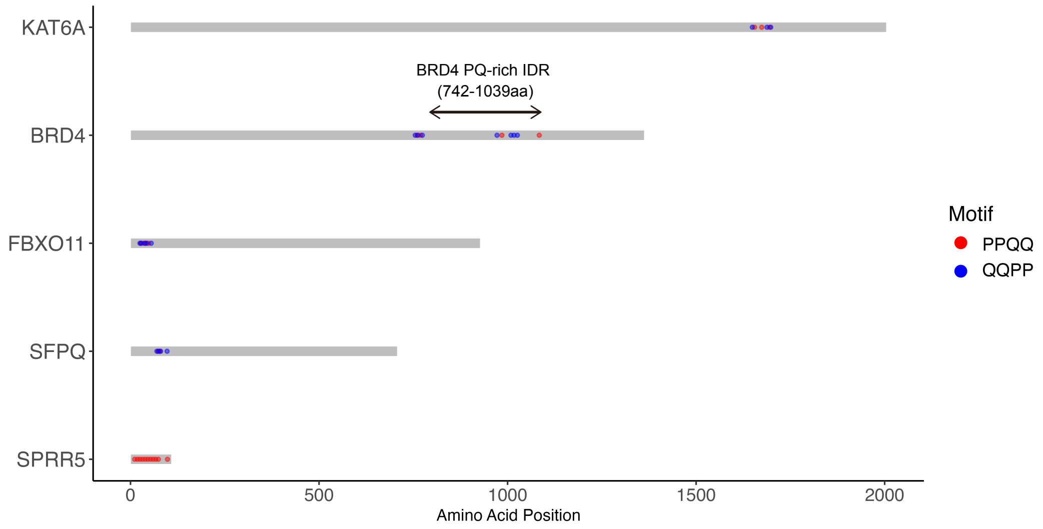


D

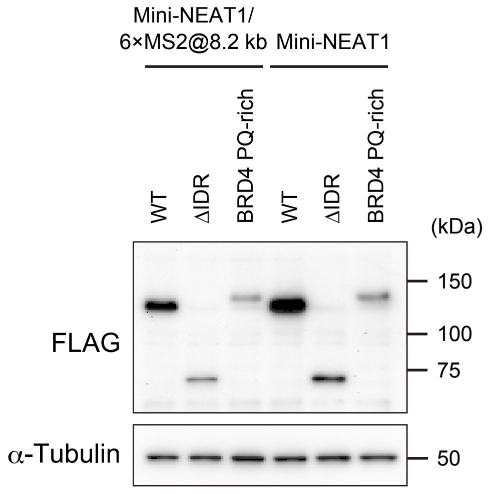

Table S1

| Reagent or Resource | Source | Identifier |
|---|---|---|
| **Antibodies** | | |
| anti-DDDDK rabbit polyclonal Ab | MBL | PM020 |
| anti-SRRM2 mouse monoclonal Ab | Sigma | S4045 |
| anti-FLAG mouse monoclonal Ab | Sigma | F3165 |
| anti-α-Tubulin mouse monoclonal Ab | Abcam | ab7291 |
| Goat anti-rabbit Alexa Fluor 405 | Invitrogen | A31556 |
| Goat anti-mouse Alexa Fluor 488 | Invitrogen | A28175 |
| Mouse IgG HRP Linked Whole Ab | Cytiva | NA931 |
| **Bacterial and Virus Strains** | | |
| DH5α Competent Cells | Thermo Fisher Scientific | 18265017 |
| Stbl3 Competent Cells | Thermo Fisher Scientific | C737303 |
| **Chemicals, peptides, and recombinant proteins** | | |
| Z-Leu-Leu-Leu-al (MG132) | Sigma | C2211 |
| Blocking reagent | Roche | 11096176001 |
| TransIT LT-1 Reagent | Mirus | MIR2300 |
| cOmplete, EDTA free Protease Inhibitor Cocktail | Roche | 5056489001 |
| ProLong Diamond Antifade Mountant | Invitrogen | P36970 |
| **Critical Commercial Assays** | | |
| Stellaris FISH probe (Human NEAT1_5 with Quaser 570 Dye) | LGC Biosearch Technologies | Custom (see Table S2) |
| Stellaris FISH probe (Human NEAT1_14.5–16.6 kb with Quaser 570 Dye) | LGC Biosearch Technologies | Custom (see Table S2) |
| n-Fusion® HD Cloning Kit | Takara Bio | 639649 |
| **Experimental Models: Cell Lines** | | |
| Human HAP1 cell line | Horizon Discovery | C631, RRID:CVCL_Y019 |
| HAP1 Mini-NEAT1 (Δ1–8 kb/16.6–22.6 kb) | Takakuwa et al., 2023 | N/A |
| HAP1 Mini-NEAT1/6×MS2@8.2 kb | Takakuwa et al., 2023 | N/A |
| **Recombinant DNA** | | |
| Plasmid: pcDNA5/FRT/TO/i/FLAG/MCP/SFPQ WT | Takakuwa et al., 2023 | N/A |
| Plasmid: pcDNA5/FRT/TO/i/FLAG/MCP/SFPQ ΔRRM2/NOPS | Takakuwa et al., 2023 | N/A |
| Plasmid: pcDNA5/FRT/TO/i/FLAG/MCP/SFPQ ΔIDR (Δ33–265aa) | Takakuwa et al., 2023 | N/A |
| Plasmid: pcDNA5/FRT/TO/i/FLAG/MCP/SFPQ Δ33–104aa | This study | N/A |
| Plasmid: pcDNA5/FRT/TO/i/FLAG/MCP/SFPQ Δ105–188aa | This study | N/A |
| Plasmid: pcDNA5/FRT/TO/i/FLAG/MCP/SFPQ Δ189–265aa | This study | N/A |
| Plasmid: pcDNA5/FRT/TO/i/FLAG/MCP/humanSFPQ zebrafishPLD | This study | N/A |
| Plasmid: pcDNA5/FRT/TO/i/FLAG/MCP/SFPQ PQPQ (33–104aa) | This study | N/A |
| Plasmid: pcDNA5/FRT/TO/i/FLAG/MCP/SFPQ PPQQ (33–104aa) | This study | N/A |
| Plasmid: pcDNA5/FRT/TO/i/FLAG/MCP/SFPQ P3Q3 (33–104aa) | This study | N/A |
| Plasmid: pcDNA5/FRT/TO/i/FLAG/MCP/SFPQ P12Q12 (33–104aa) | This study | N/A |
| Plasmid: pcDNA5/FRT/TO/i/FLAG/MCP/SFPQ P36Q36 (33–104aa) | This study | N/A |
| Plasmid: pcDNA5/FRT/TO/i/FLAG/MCP/SFPQ P3Q3×3 (33–188aa) | This study | N/A |
| Plasmid: pcDNA5/FRT/TO/i/FLAG/MCP/SFPQ P3Q3×8 (33–188aa) | This study | N/A |
| Plasmid: pcDNA5/FRT/TO/i/FLAG/MCP/SFPQ P3Q3×12 (33–188aa) | This study | N/A |
| Plasmid: pcDNA5/FRT/TO/i/FLAG/MCP/SFPQ BRD4 PQ-rich | This study | N/A |
| **Software and Algorithms** | | |
| R | CRAN | https://cran.r-project.org/ |
| R studio | Posit | https://posit.co/download/rstudio-desktop/ |
| CellProfiler | BROAD INSTITUTE | https://cellprofiler.org/ |
| Python | Python Software Foundation | https://www.python.org/ |
| g:Profiler | BIIT | https://biit.cs.ut.ee/gprofiler/gost |
| GENESIS | RIKEN | https://www.r-ccs.riken.jp/labs/cbrt/ |
| ImageJ/FIJI | NIH | https://imagej.nih.gov/ij/ |
| NARDINI+ | GitHub/Google Colab Notebook | https://github.com/mshinn23/nardini |

Table S2

| Human NEAT1_5' probe set | Human NEAT1_14.5-16.6 kb probe set |
|---|---|
| caagttgaagattagccctc | ctcacaaaatccaggaacca |
| agcccttggtctggaaaaaa | gctggattagccttttcaaa |
| aagttcagttccacaagacc | gttaggagaaaacatggggt |
| caggccgagcgaaaattaca | aagtctagatagagtcctcc |
| ctgtcaaacatgctaggtgc | cctgtcacttgttatgctaa |
| aagcgttggtcaatgttgtc | ggtgtaaggacaacaggcta |
| gtggagtgagctcacaagaa | gttaatagctgtattagcca |
| cttaccagatgaccaggtaa | aaccattaggaactggcacg |
| ttaccaacaataccgactcc | tcgctattttgaagtgcaga |
| cggtccatgaagcatttttg | aataatgaggtagggcttcc |
| tcgccatgaggaacactata | tagccaggagaattctggaa |
| atctgcaggcatcaattgag | aaggtaaacatgcagcctgg |
| agcaaggcctggaaacagaa | cggcatgaagacatcacagg |
| catctgctgtggacttttta | gacgtaccttattcttgcat |
| ttcatgggctctggaacaag | aatagacgtgagtggatgga |
| gatgcagcatctgaaaacct | gctttctgtgtaatgcaaag |
| aaactagtatgaccggaggc | cgaggtagacagaccaagac |
| ttgaagcaaggttccaagca | ggcagtgaggacaactagat |
| tgttctacagcttagggatc | aagtactctgtgatggggta |
| tacaaggcatcaatctgcgt | gtaaccattaggcagagcaa |
| caaacaggtgggtaggtgag | gcaagctgggtttgtatgaa |
| cttctccgagaaacgcacaa | ctacgctctctgaatgtgac |
| ccaagttatttcatcaggct | tggaaaacaacccatcccag |
| tctaatatatccccagtcta | gaagagctagccacacagtg |
| cacaacacaatgacacccctt | acattccgggtacacagaac |
| caaactagacctgccatttc | gctggacactagaacaggac |
| ctcctagtaatctgcaatgc | agcaggaataggctgttgag |
| aaagagcactaccggtgtac | aaaaagttccagcaggctgt |
| tcctcttactagaatgccaa | ccagcatggcaacatatttt |
| ctaagcaacttctcacttcc | tgaaaaaggctgccaggggg |
| taacacttcttcagtcttcc | gcacagccaaatcaagtcac |
| cctttggttctcggaaaact | |
| tgtgagatggcatcacacac | |
| ccaggaggaagctggtaaag | |
| ctctgaaacaggctgtcttg | |
| tcacttgataacacccacac | |
| cagcgaaggatgctgatctg | |
| atcaaccacctaagttgcta | |
| gtggtcccttaaatacgtta | |
| agaagagcccatctaatctc | |
| gatgtgtttctaaggcacga | |
| ggtcttgttttccaaactga | |
| catgtagtaaaggcacctcg | |
| ccattggtattactttacca | |
| ctctaaatcccaacgacagt | |
| atttcacaacagcatacccg | |
| ccagtactttcaaccatcta | |
| agttcttaccatacagagca | |
